# Ocular elongation and retraction in foveated reptiles

**DOI:** 10.1101/2021.01.20.427408

**Authors:** Ashley M. Rasys, Shana H. Pau, Katherine E. Irwin, Sherry Luo, Paul A. Trainor, Douglas B. Menke, James D. Lauderdale

## Abstract

**Background:** Pronounced asymmetric changes in ocular globe size during eye development have been observed in a number of species ranging from humans to lizards. In contrast, largely symmetric changes in globe size have been described for other species like rodents. We propose that asymmetric changes in the three-dimensional structure of the developing eye correlate with the types of retinal remodeling needed to produce areas of high photoreceptor density. To test this idea, we systematically examined three-dimensional aspects of globe size as a function of eye development in the bifoveated brown anole, *Anolis sagrei*.

**Results:** During embryonic development, the anole eye undergoes dynamic changes in ocular shape. Initially spherical, the eye elongates in the presumptive foveal regions of the retina and then proceeds through a period of retraction that returns the eye to its spherical shape. During this period of retraction, pit formation and photoreceptor cell packing are observed. We found a similar pattern of elongation and retraction associated with the single fovea of the veiled chameleon, *Chamaeleo calyptratus*.

**Conclusions:** These results, together with those reported for other foveated species, support the idea that areas of high photoreceptor packing occur in regions where the ocular globe asymmetrically elongates and retracts during development.

**Key Findings:**

- The eyes of the brown anole, *Anolis sagrei*, and veiled chameleon, *Chamaeleo calyptratus* undergo dynamic asymmetrical changes in ocular shape during development.
- In both species, asymmetric elongation and retraction of the ocular globe is associated with fovea morphogenesis.
- Pit formation and photoreceptor cell packing in the foveal area occur when the corresponding region of the ocular globe is retracting relative to adjacent regions.

## Introduction

Decades of experimental work have revealed a great deal about the developmental mechanisms that govern the patterning, differentiation, and growth of the vertebrate eye. Much of this understanding has come from functional studies in mouse, rat, chicken, *Xenopus*, zebrafish, and medaka.^1,2^ Additional insights have come through genetic studies of human syndromes that feature eye defects.^3^ Consequently, we know a great deal about the genes and signaling pathways that regulate development of the core structures common to all vertebrate eyes, including the cornea, lens and retina.^4-7^ Missing from our current understanding of vertebrate eye development, however, is detailed knowledge about the developmental pathways that regulate the formation of specialized structures that are only present in the eyes of certain vertebrate species.

These structures include the conus papillaris, areas or streaks, and foveae. Studies of eye formation in diverse vertebrate groups are needed to determine how these specialized structures form and to achieve a more complete understanding of vertebrate eye development and evolution. Notably, modern investigations of eye development have almost completely excluded reptiles, a tremendously successful amniote group represented by over 10,000 extant species.^8^

Although eye development in reptiles remains poorly studied, other aspects of reptilian biology have been actively explored. For instance, *Anolis*, a lizard genus with approximately 400 recognized species, has served as an important model system for studies of evolution, ecology, physiology, behavior, and neuroendocrinology for many years.^9^ More recently, *Anolis* has also emerged as a system to investigate reptile development and the mechanisms that contribute to morphological evolution.^10-16^ The brown anole lizard, *Anolis sagrei*, is particularly well-suited for developmental studies due its small size, ease of husbandry, continuous egg production, high fertility, and low cost. In addition, *ex ovo* culture systems and gene-editing have been established for this species, which presents the opportunity for pharmacological and genetic manipulation of *Anolis* embryos during development.^12,17,18^ Of interest for studies of eye development, *Anolis* lizards possess specialized structures that include a bifoveated retina and a highly vascularized conus papillaris.^19,20^

Here we describe morphological and histological aspects of eye development in *A. sagrei*. We pay particular attention to alterations in ocular globe shape, which is an interesting, but poorly understood, aspect of eye development. Although typically only studied postnatally in the context of myopia in humans,^21-23^ changes in ocular shape during embryonic development have been observed in a number of foveated species, including humans,^24-30^ and non-human primates,^26,31^ as well as geckos,^32,33^ suggesting the presence of a conserved morphogenetic mechanism. The bifoveated brown anole is a good model system in which to study the mechanisms underlying fovea development in a vertebrate eye. In this study, we provide the first systematic three-dimensional assessment of the dynamic changes in ocular shape with an emphasis on ocular elongation and retraction and its association with fovea formation.

## Material and methods

### Animals

All experimental procedures were conducted in accordance with the National Institutes of Health Guide for the Care and Use of Laboratory Animals under protocols approved and overseen by the University of Georgia (anoles) and Stowers Institute for Medical Research (chameleons) Institutional Animal Care and Use Committees. *Anolis sagrei* lizards were maintained in a breeding colony at the University of Georgia following guidelines described by *Sanger et al*., 2008.^34^ Eggs were collected weekly from natural matings and placed in 100 × 15 mm lidded petri dishes containing moist vermiculite and incubated at 27-28°C and 70% humidity. Adults and hatchlings were euthanized using methods consistent with the American Veterinary Medical Association (AVMA) Guidelines for the Euthanasia of Animals.^35,36^ *Chamaeleo calyptratus* were maintained in a breeding colony at the Stowers Institute for Medical Research (Kansas City, Missouri) following guidelines described by Diaz et al., 2015^37^ and Diaz et al., 2019.^38^ Eggs were collected at the time of oviposition and incubated at 26-28°C and 50% humidity on damp vermiculite. Male and female embryos of both species were used for these studies.

### Staging

Embryonic development of *Anolis* lizards typically takes place over a 30-33 day period, starting with fertilization, which takes place internally.^39^ Early embryogenesis proceeds within the oviduct. *A. sagrei* embryos obtained from eggs that were collected after egg-laying were staged as described by Sanger et al., 2008.^39^ Embryos younger than those captured by the Sanger staging series were denoted with the prefix “PL” for pre-laying followed by a number. We describe here 5 PL timepoints, which includes the first few embryos of the Sanger staging series (Sanger St 1-3 correspond to PL 3-5). PL stage embryos were collected from gravid adult females following euthanasia. *C. calyptratus* embryos were staged following criteria described by Diaz et al., 2017 and Diaz et al., 2019 and stage matched to the anole using Sanger’s morphological criteria.^38-40^

### Dissection

Lizard embryos were removed from their shells using a blunt pair of forceps and iris scissors in 1x phosphate-buffered saline (PBS; 137 mM NaCl, 2.7 mM KCl, 10 mM Na_2_HPO_4_, 1.8 mM KH_2_PO_4_, pH 7.4). Upon removal of yolk and amniotic sac with fine forceps, embryos were placed into 60 ml of fresh 1x PBS solution with 1 ml of 0.4% pharmaceutical grade, neutrally buffered tricaine (TRICAINE-S; Western Chemical Inc) to anesthetize for imaging. Eyes were enucleated from embryo stages >4 with fine forceps and placed in Bouin’s fixative at 4°C overnight on a rocker. Following fixation, eyes were washed five times at 15 min per wash in 1x PBS. Specimens were stored in 70% ethanol solution (EtOH) at 4°C until processed for histology. Whole chameleon embryos were dissected from eggs in a similar manner, fixed in Bouin’s at 4°C overnight, washed in 1x PBS, and stored in 70% EtOH prior to shipment from the Stowers Institute for Medical Research. Upon arrival at the University of Georgia, embryos were slowly rehydrated in a series of graded EtOH/PBS solutions. Once fully rehydrated, eyes were carefully removed from embryos.

### Whole Eye Measurements

Prior to fixing, anole eyes were positioned in both a lateral and dorsal orientation and imaged with a ZEISS Discovery V12 SteREO microscope and AxioCam while in 1x PBS solution. AxioVision 4 software (Carl Zeiss MicroImaging) was used to take axial measurements along the dorsoventral (y), nasotemporal (x), and lateromedial (z) aspects of imaged eyes from embryos at Sanger stages 4-18, the hatchling (Hch), and the adult (Adt). The lateral view, which encompassed the whole cornea, was used to measure the y- and x-axes, whereas the dorsal view was imaged to obtain x- and z-axes. Because x-axial measurements can be acquired from both views, we included only the dorsal x-axial value in our dataset and used the other value as a control for proper orientation. Measurements from eyes that were not correctly positioned were excluded from the dataset. We normalized the x- and z-axial measurements from each lizard to that same individual’s y-axis and then multiplied this number to the mean (µ) of that individual’s stage group y-axis (µ_y_), 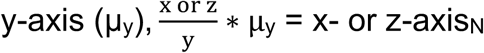. To identify trends in ocular growth, a ratio was then calculated for each axis by taking the raw y-axis dataset and the normalized datasets (x- and z-axis_N_) from every lizard and dividing these values with the corresponding µ of the raw y-, x-, and z-axial lengths of the hatchling, 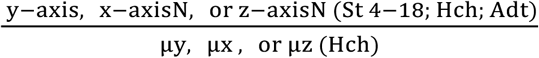 Chameleon eyes were processed as reported above except imaging was performed post-fixation and rehydration. Prism 7 (GraphPad Software) and JMP V14.1 (JMP SAS) were used for graph generation and data analyses. As a few of the sample groups did not have a normal distribution, we used a nonparametric one-way ANOVA (Kruskal-Wallis) and Mann-Whitney for our statistical analyses.

### Paraffin Sectioning

Eyes were dehydrated in a series of graded ethanol solutions 70%, 80%, 90%, 96% and 100% (twice) for a minimum of 15 min each and then soaked in xylene for a total of 30 min for all embryonic stages. Tissue specimens were incubated in a series of 3 paraffin wax jars for 30 min at 65°C, embedded in paraffin, and serially sectioned horizontally at 10 μm. In adult specimens, the dorsal aspect of the eye was punctured with a 0.15 mm minutien pin prior to processing in xylenes and paraffin waxes to facilitate wax entry. Processing time in xylene in the adult was extended up to a total of 2 hrs. Eyes were serial sectioned on a horizontal plane. Sections were stained with hematoxylin & eosin following standard protocols and mounted in Cytoseal (Thermo Scientific™ Richard-Allan Scientific™). Photomosaic images were generated using a KEYENCE BZ-700 microscope with Keyence image stitching software. Adobe Photoshop CC (2017.01 release) was used to digitally enhance contrast and adjust white balance of images.

## Results

### Anatomy of the adult anole eye

Laterally positioned in the skull, the adult eye externally is oblate spheroid in shape with a prominent convex cornea slightly biased toward the nasal region (Figure 1a,b). Peripheral to the cornea is the sclera sulcus, whose curvature is supported by 14 sclera ossicles (Figure 1b,c). The sclera ossicles are uniformly orientated in a patterned ring except for a slight extension in the temporal region of the eye (Figure 1c). The anole’s radial pupil is positioned centrally, fashioned by a heavily pigmented iris with an array of iridophores and melanophores that extend into the circumferential sclera sulcus (Figure 1b,c). The iris is asymmetric with dorsal and ventral notches defining the boundary between the larger temporal and smaller nasal region (Figure 1c, arrowheads). Dorsally, a protruding blood vessel is present that extends from the optic nerve, wraps around the region of the center fovea (a small bulge in the medial region), and dissipates towards the dorsal nasal area of the eye (Figure 1b, narrow arrowheads). The optic nerve (not shown) exits the eye ventrally and temporally to the central fovea.

**Figure 1.**
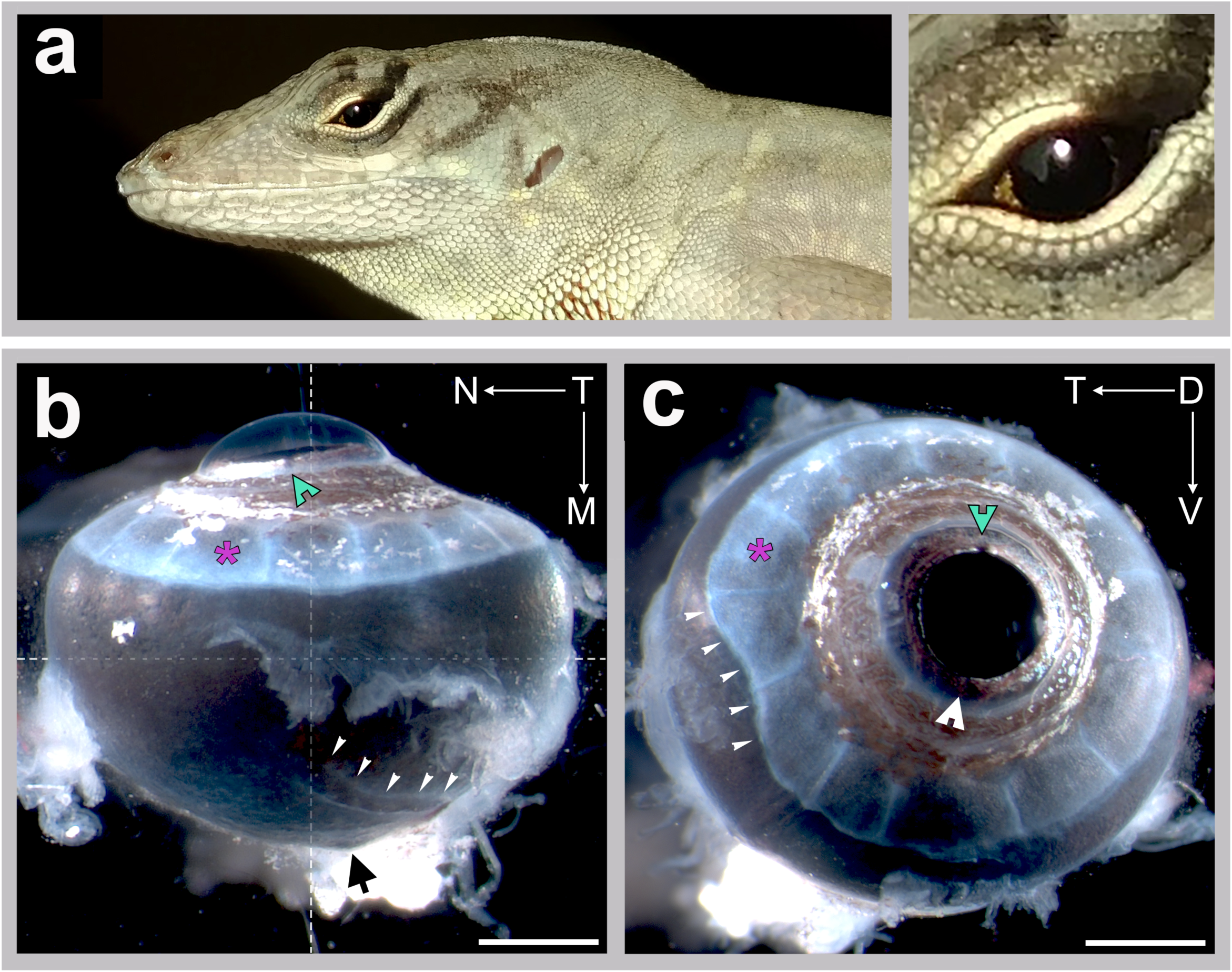
Adult anole eye. Top panel (a) shows adult male lizard with an enlarged view of its left eye. Bottom panels show a lateral view (b) and dorsal view (c) of a right eye. Directionality is designated by arrows and the letters T – temporal, N – nasal, D – dorsal, and V – ventral. Markers indicate: white arrow heads with notches – dorsal (green) and ventral (white) iris notches; asterisks (magenta)– individual sclera ossicle sheets; narrow white arrow heads (b) – dorsal blood vessel, (c) temporal sclera ossicle deformation; black arrow – optic nerve; crosshair – dorsal optical axis; and scale bars – 1 mm.

Internally, the adult anole eye is very similar to other vertebrates, possessing a cornea, iris/ciliary body, lens, and retina (Figure 2a). Anteriorly, the transparent cornea, which is exterior to the lens and iris, is composed of epithelial, stromal, and endothelial layers (Figure 2a, Rasys et al., *in prep*). These layers thicken towards the limbal area where the cornea and anterior margin of the sclera ossicles meet. The sclera ossicles extend from the limbal area outward towards the sclera proper in overlapping sheets making up the sclera sulcus. Underlying this sclera sulcus is the long thin ciliary body (Figure 2a). The ciliary body lacks a ciliary process and comprises an inner non-pigmented layer and an outer pigmented epithelial layer (Rasys et al., *in prep*). These layers extend from the neural retina to the iris-ciliary boundary. Beyond this boundary is the iris proper which is closely associated with the lens (Figure 2a). Both the inner and outer iris epithelium are pigmented. The lens is oval shaped with a central nucleus, inner cortex, and outer annular pad (Figure 2a). Anchoring the lens are zonular fibers that extend from the ciliary inner non-pigmented epithelium (Figure 2a). Posteriorly, the eye includes the retina, retinal pigmented epithelium, choroidal, and sclera layers. The neural retina is avascular and composed of a ganglion cell layer (GCL), inner and outer plexiform layers (IPL and OPL), inner and outer nuclear layers (INL and ONL) (Figure 2b,c).

**Figure 2.**
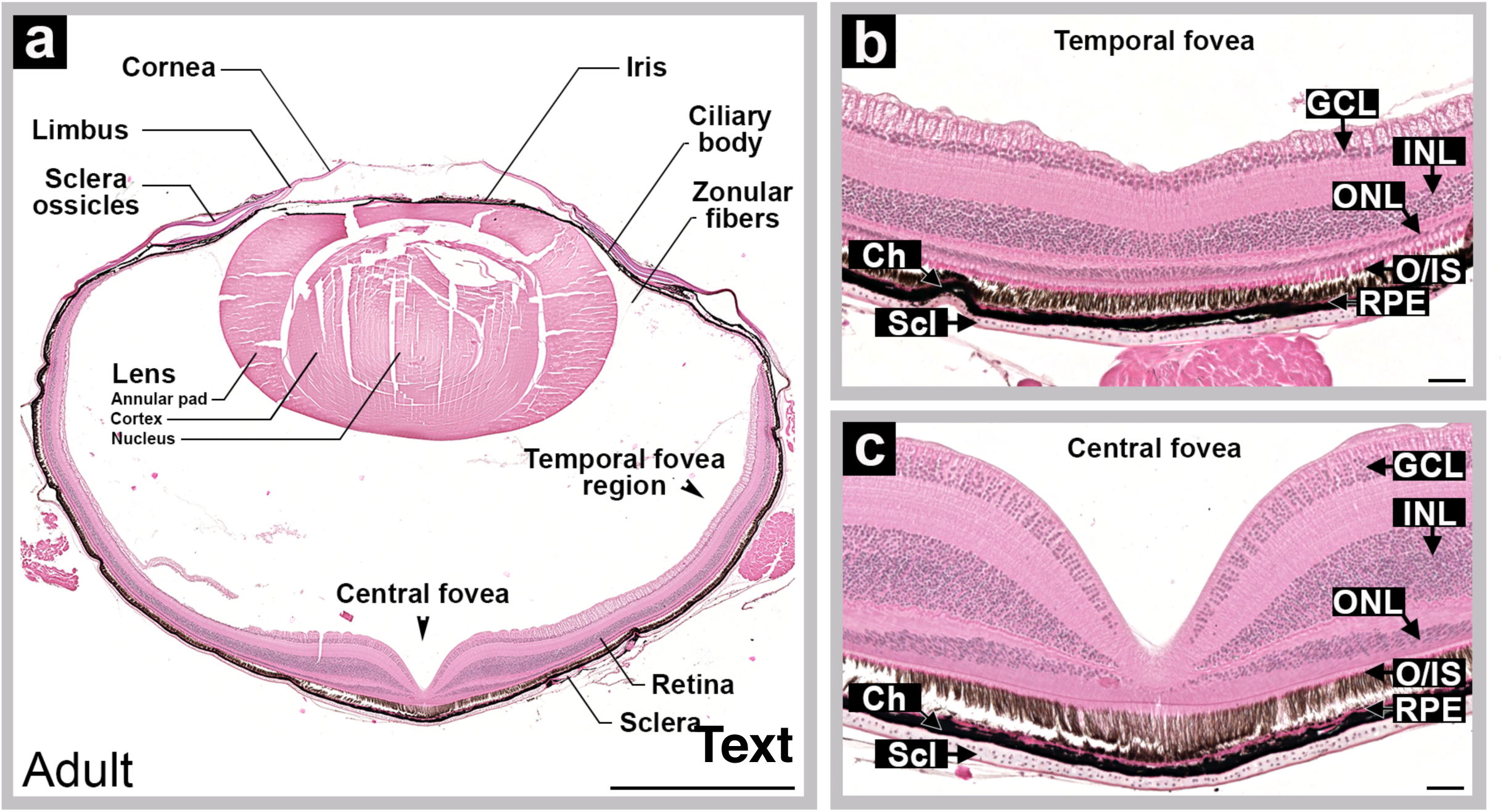
Anole eye organization and retinal layout. Left panel (a) shows a diagram labeling structures present in a horizontally sectioned adult eye. Right panels show magnified views of temporal (b) and center (c) foveae. Note: temporal fovea shown in (b) is 150 µm deep to the plane of section in (a). Markers indicate GCL – ganglion cell layer; INL – inner nuclear layer; ONL – outer nuclear layer; RPE – retina pigmented epithelium; Ch – choroid; Scl – sclera; and scale bars – 1mm (a) and 50µm (b and c).

Each eye has two foveae with one in the central retina and a second in the temporal retina (Figure 2). The central fovea, located slightly temporal (4-5 degrees) to the optical axis (the central or mid-point of the eye), exhibits a distinct pit accompanied by a higher density of photoreceptor cells compared to the peripheral retina, as has been previously described for other anoles.^19,20^ The retina at the central foveola is devoid of the GCL, INL and ONL (Figure 2a,c). The retina in the parafoveal region has a larger number of cell bodies in the INL and ONL compared to the peripheral retina (Figure 2a,c). A second, shallower fovea is located in the temporal retina roughly 45 degrees from the optical axis or 40 degrees from the visual axis (central fovea) (Figure 2). This temporal fovea exhibits a shallow pit accompanied by an increase in photoreceptor cell density. At its center, all the retinal cell layers are retained, but layers are thinner than the surrounding peripheral retina (Figure 2b). The temporal fovea is approximately one-third the area of the central fovea, with fewer photoreceptor cells.

### Eye formation

In anoles, fertilization takes place internally and embryonic development occurs over a 30-33 day period.^39^ As a consequence, the earliest stages of eye development take place while the eggs are in the oviduct. Embryonic stages pre-egg lay are denoted with the prefix “PL”. Stages post-egg lay use the Sanger staging series^39^ and are denoted using the prefix “St”. Stages PL 3-5 correspond to Sanger stages St 1-3.^39^ For these earliest timepoints we use the PL nomenclature to help distinguish pre-lay stages (PL 1-5) from post-lay stages (Sanger St 4-19).

The initial stages of eye development occur between PL 1 and PL 5. Optic vesicle formation is evident in embryos at PL 1 (Figure 3a), which is about 1 day after fertilization and 3 days before egg lay. The lens placode is visible at PL 2 (see developmental poster in supplementary data). By PL 3, the optic cup and lens pit are present (Figure 3b). Rapidly following this period, the cornea separates (PL 5) from the newly minted lens vesicle (PL 4) and the optic fissure begins to close (PL 5) (developmental poster). At the time of egg lay (∼St 4), the anole eye is spherical, trace amounts of pigmentation are evident in the temporal region of the eye (Figure 3c) and present nasally by St 5 (Figure 3d). At this time (St 5), the prospective central and temporal foveal regions are evident as thickenings in the retina, also defined as retinal mounding (Figure 3g, *Rasys et. al*., in prep). St 5 also marks the first appearance of the sclera sulcus, evidenced by a slight depression in the temporal area, which extends to the nasal side by St 6 (Figure 3d). Eventually, the sclera sulcus forms a complete ring encircling the cornea (developmental poster, *whole eye stages 6-8*). Shortly following this period, pigment begins to increase - initially in the iris between St 6-10 and then throughout the rest of the eye. In the iris, pigment is deposited first as a narrow band along the horizontal axis (St 6-7) before radiating outward throughout the dorsal and ventral regions (St 8-9) (Poster, *Whole eye*). By St 10, pigment in the iris is black and evenly distributed. Granules are also just becoming obvious throughout the whole eye, but more so in the temporal region. At St 13, retinal mounding is no longer present in the central and reduced in the temporal foveal regions (Figure 3h). The eye is a light brown color which darkens between St 15-17 and is completely black by the time of hatching (Figure 4b; developmental poster). During this period the sclera ossicles that shape the sulcus and provide support to the underlying ocular structures, are starting to form. The sclera anlagen first manifests as a ring of pale conjunctiva papillae around the cornea between St 11-12. By St 13, scleral sheets are present although they are small and by St 15-16 these expand, radiating outward and eventually overlapping (St 17) with the neighboring plates (developmental poster). Iridophores (reflective pigments) scattered throughout the iris and sclera sulcus region, are also apparent during this time.

**Figure 3.**
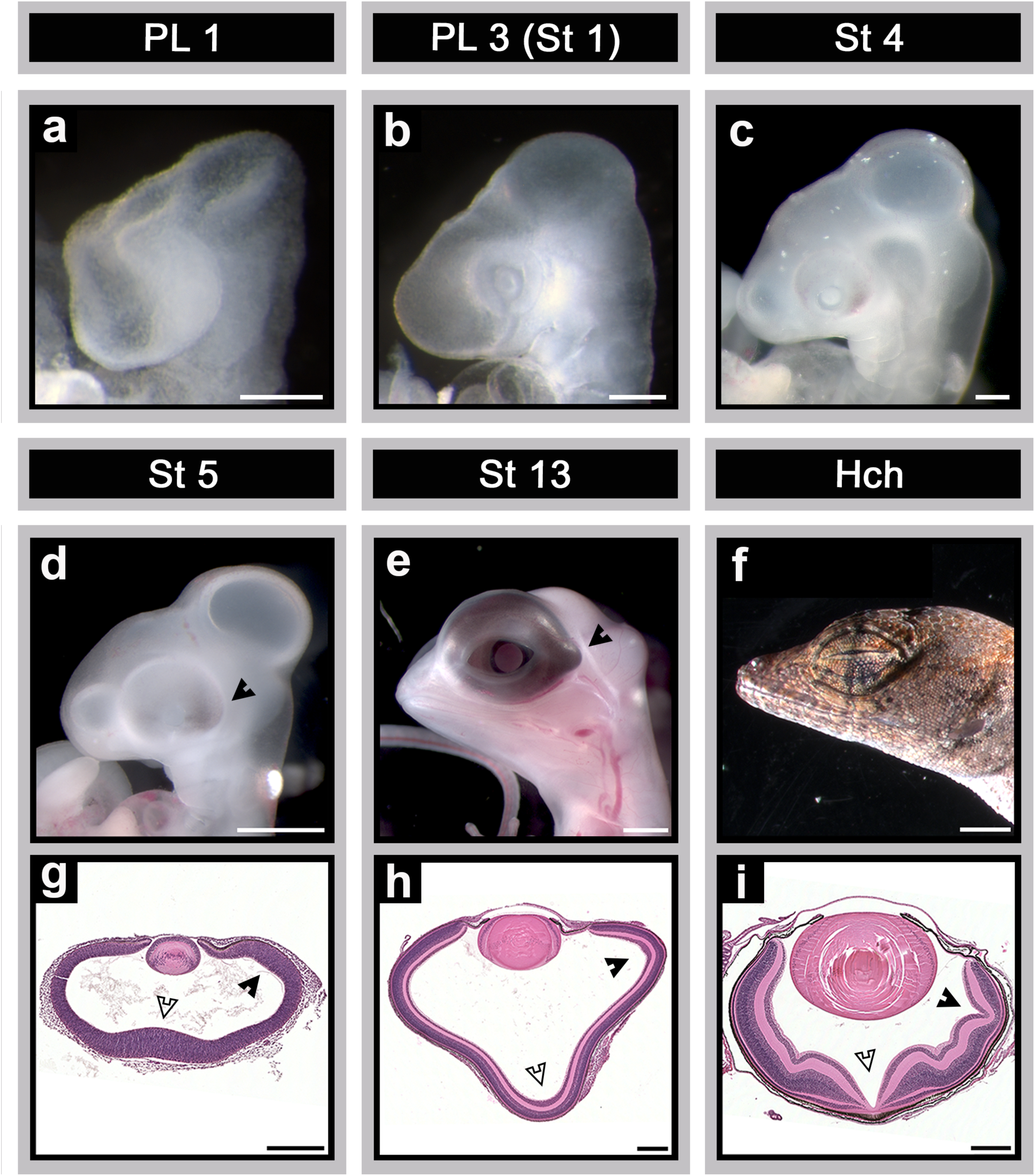
Developmental series of early, mid, and late embryonic stages of eye morphogenesis. Top panel shows an array of early-stage embryos (PL1, PL3, and St 4) during optic vesicle (a), len’s pit (b), and lens separation (c). Bottom panel shows later stages (St 5, 13, and Hch) along with histological sections through the eye’s center horizontal plane. Black arrow – temporal eye region; open arrows – center and closed arrows – temporal retinal and fovea regions; and scale bars are 250 µm (a-c, g-i) and 1 mm (d-f).

**Figure 4.**
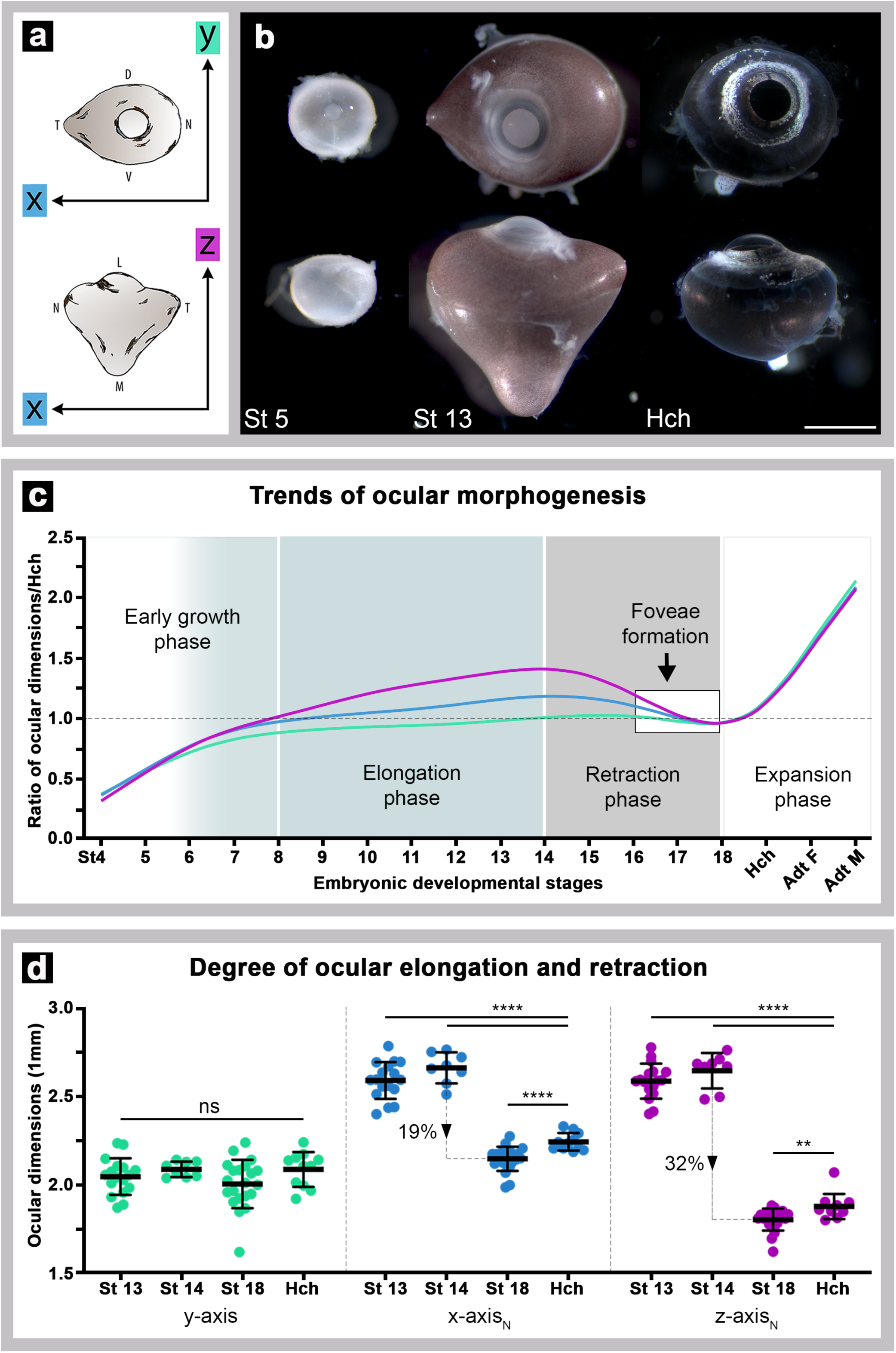
Anole eyes undergo asymmetrical ocular elongation followed by retraction during embryonic development. (a) Diagram illustrating orientation of y- (green), x- (blue), and z-axis (magenta) ocular measurements. (b) Lateral (top) and dorsal (bottom) views of whole right eyes from early (St 5), mid (St 13), and late (Hch) stages. Ocular elongation and retraction phases are evident in the stage 13 embryo and hatchling, respectively. All eyes are to scale with one another; scale bar is 1 mm. (c) Graph displaying trends in ocular morphogenesis throughout development (St 4-Hch) and adulthood (Adt F – adult female; Adt M – adult male). This graph was generated by taking each lizard’s ocular dimensions (y-axis and normalized x- and z-axis_N_) and dividing it by the mean (µ) hatchling ocular dimensions (y-, x-, and z-axis) to calculate a ratio. (d) Direct comparison of ocular length along different axes reveals the degree of ocular elongation (St 13-14) and subsequent retraction (St 18-Hch) in the anole eye.

### Eye morphogenesis

The anole eye exhibits conspicuous asymmetric changes in morphology during development (Fig 4). To assess potentially complex changes in the three-dimensional shape of the globe, measurements were made along the three anatomical axes of the eye at different stages of development. For this study, the dorsoventral axis was defined as the y-axis, the nasotemporal axis was defined as the x-axis, and the lateromedial axis was defined as the z-axis (Figure 4a). The lateromedial (z) axis is also the optical axis and passes through anterior (iris and lens) and posterior (retina) structures of the eye.

At St 5, the ocular globe is mostly spherical in shape (Figure 4b), with similar lengths along the three anatomical axes. At late mid-gestation (St 13-14), the ocular globe has a triangular appearance with increased lengths along the nasotemporal (x) and lateromedial (z) axes compared to the length of the dorsoventral axis (Figure 3e,h; Figure 4b). Interestingly, whereas the nasal surface of the globe has a rounded appearance, both the temporal and medial surfaces have angular shapes. These differences in surface geometry suggest that elongation in the nasotemporal and lateromedial planes occurred largely by changes in the temporal and medial regions of the globe, respectively. The medial region of the globe corresponds to the area of the retina that gives rise to the central fovea, and the temporal region corresponds to that of the temporal fovea (Figure 3g-i). The eye at this stage also exhibits considerably less pigmentation at the medial surface compared to the lateral surface (Figure 4b). At hatching, the ocular globe has a spherical shape with uniform pigmentation (Figure 3i; Figure 4b). Remarkably, the globe at this stage has a smaller surface area than that of mid-gestational embryos (Figure 4b). The change in morphology and size of the globe in a hatchling compared to that of a mid-gestational embryo suggests that the eye undergoes asymmetric retraction in the areas encompassing the developing central and temporal foveae during the period between mid-gestation and hatching.

To better understand the dynamics of globe morphogenesis in the brown anole, axial measurements along the three anatomical planes were made of eyes at embryonic stages St 4-18, hatchlings (Hch), and adults (Adt) (Figure S1; Table S1). To facilitate a quantitative comparison of changes in the three-dimensional shape of the eye, the three measurements of each eye were converted to a standardized metric (see *Whole Eye Measurements* in Methods). Briefly, in this metric the axial length of the y-axis, which did not exhibit appreciable elongation and retraction during development, was used as a normalization factor for measurements along the x- and z-axes. Variance due to differences in embryo body and eye size within an individual’s stage group was handled by multiplying the normalized measurements for each individual by the group mean value (µ) for the y-axis (Figure S1; Table S1). This operation resulted in an overall reduction in variance seen at each stage in both x- and z-axis_N_ datasets, which suggests that the degree to which an eye elongates and retracts is proportional to the embryo’s eye size (Figure S2).

Using the hatchling eye as a reference, regional differences in globe morphogenesis as a function of developmental stage were assessed by taking the raw y-axis dataset and the normalized datasets (x- and z-axis_N_) from every lizard and dividing these values with the corresponding mean of the raw y-, x-, and z-axial lengths of the hatchling (see *Whole Eye Measurements* in Methods). This analysis revealed four distinct phases in ocular morphogenesis (Figure 4c).

Phase 1, which occurs between embryonic St 4-8, is characterized by rapid growth of the eye. In embryos at St 4-5, the globe appears to expand uniformly along all 3 axes, which suggests that ocular growth during this period is equally distributed across the eye. Although the St 5 eye is 50% smaller than the hatchling eye, the eye at these two stages is quite similar in overall shape (Figure 4b,c; Table S1). At St 6, the globe begins to exhibit asymmetric expansion, with more growth along the nasotemporal and lateromedial axes compared the dorsoventral axis. By St 7, expansion along the lateromedial axis is greater than along the nasotemporal axis.

The start of the phase 2 is defined as the developmental timepoint when morphological asymmetry of the globe is clearly visible. This condition is met at St 8. It is also at this point that the overall globe size is similar to that of the hatchling. Between St 8 and the close of the second phase at St 14, the globe continues to expand asymmetrically along the nasotemporal and lateromedial axes, with the more pronounced expansion along the lateromedial axis (Figure 4c,d). The globe reaches maximum lengths along the nasotemporal and lateromedial axes by St 14. At this stage the length of the lateromedial axis is 1.4x that of the hatchling, and the length of the nasotemporal axis is 1.2x that of the hatchling (Figure 4b,c).

Phase 3 is the epoch during which the globe begins to shorten in length along the nasotemporal and lateromedial axes to regain a spherical shape. This phase encompasses St 15-18 (Figure 4c,d), and corresponds to the time during which the fovea acquires its distinctive morphological characteristics (Figure 3h,i). Upon completion of retraction, the eyes are slightly smaller than those of hatchlings (St 17-18; Figure 4c,d). The fourth, and final, phase is characterized by a uniform expansion of the globe, which begins at the close of St 18 and continues into adulthood (Figure 4c). From hatching, the eye doubles in size by the time the lizard reaches adulthood (Figure 4c).

To determine the magnitude of the asymmetric shape changes during ocular morphogenesis, measurements capturing the maximum extent of elongation (St 13-14, Figure 4c) were compared to those capturing the maximum extent of retraction (St 18-Hch, Figure 4c). Axial measurements of globes from lizards at St 13, 14 and 18, and hatching were compared using one-way ANOVA analyses (nonparametric Kruskal-Wallis test); normalized values were used for the nasotemporal and lateromedial axes (for mean and standard deviation, see Table. S1). Although measurements along the dorsoventral axis were not statistically different across these four developmental time periods (p-value 0.1585; alpha = 0.05), significant differences were observed for measurements along both the nasotemporal and lateromedial axes (p-value <0.0001 for both; alpha = 0.05).

Differences between the groups were compared using the Mann-Whitney test (Figure 4d). Among the normalized nasotemporal (x-axis_N_) datasets, St 13 and 14 measurements were significantly different from the St 18 and the hatchling (p-value <0.0001 for both; alpha = 0.05) but not between St 13 and 14 (p-value 0.1244). Similar results were observed among the normalized lateromedial (z-axis_N_) measurements. For St 18 compared to hatchling, x-axis_N_ and z-axis_N_ lengths were significantly different (p-value <0.0001, and p-value of 0.0017, respectively). This suggests that by St 18 ocular retraction is finished and the expansion phase is already well underway by the time of hatching. The large difference in mean x-and z-axis_N_ lengths between St 14 and 18 (z-axis_N_ 2647 to 1807 µm; x-axis_N_, 2663 to 2151 µm) equated to a 32% and 19% reduction in the central and temporal regions, respectively (Figure 4d).

Changes in intraocular pressure can be one mechanism that drives globe expansion and retraction. Using an inflated ball as a model, morphological indicators of a pressurized globe could include a taut ocular surface that resists deformation. Conversely, using an under-inflated ball as a model, a previously-pressurized globe that has lost pressure might exhibit wrinkles or folds of the surface and be more flaccid. The surfaces of globes prior to St 16 were smooth and taut to the touch. In contrast, starting at St 17, the exterior ocular surface was easily depressed with a pair of dull forceps and wrinkles could be clearly seen in about half of the eyes examined (7/13 St 17 lizards) (Figure 5). At St 18, wrinkling could still be detected (8/19 lizards), although it was not as pronounced. By hatching, the majority of globes (9/10 lizards) were completely smooth and once again the ocular surface was taut (Figure 5a). For developmental stages between St 14 and hatching, wrinkled eyes tended to have reduced normalized measurements along the lateromedial (z-axis_N_) compared to smooth eyes (Figure 5c).

**Figure 5.**
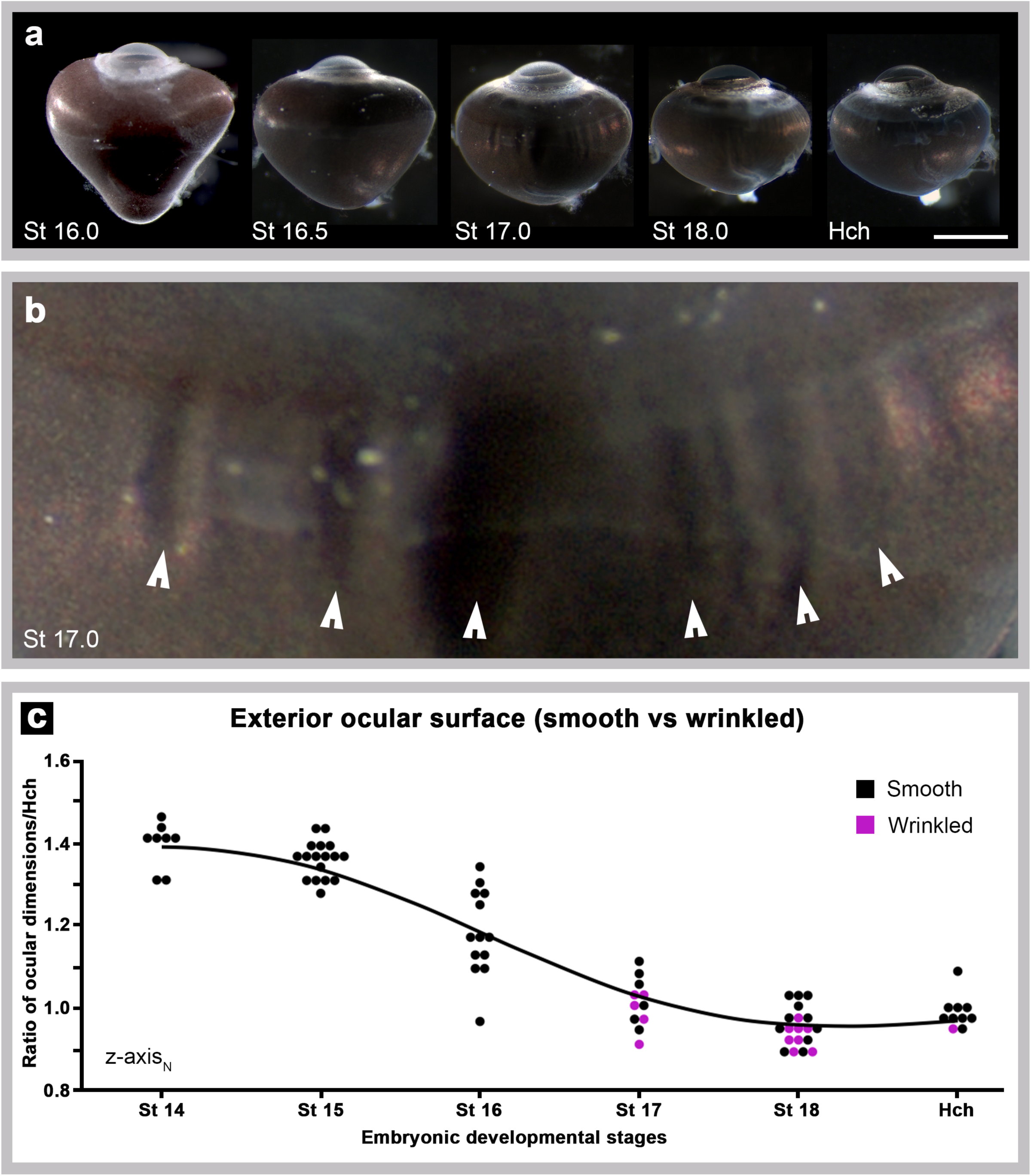
The formation of surface wrinkles coincides with ocular retraction. (a) Dorsal views of whole right eyes from stages 16 - Hatchling showing the progressive steps of ocular retraction; scale bar – 1 mm. (b) An enlarged view of the St 17 eye from panel (a); arrow heads mark folds/wrinkles present along the outer ocular surface. (c) ocular dimensions along the z-axes relative to the hatchling eye; eyes where folds/wrinkles were observed are indicated in magenta.

### Ocular elongation & retraction in other foveated lizards

To test if ocular elongation and retraction occurs in other foveated lizards, ocular morphogenesis was examined in the veiled chameleon lizard, *Chamaeleo calyptratus*. This species was chosen because it possesses a single, prominent central fovea that is completely devoid of all retina cell layers.^41^ Chameleon embryos were collected at several time points throughout development. To facilitate comparison with anoles, chameleon embryos were staged following criteria described by Diaz et al., 2017 and Diaz et al., 2019 and then matched to the anole using Sanger’s morphological criteria.^38-40^ Only embryonic stages were used for this study; chameleon embryos collected just prior to the expected hatching date are denoted as pre-hatch “pHch” (Table S2).

As observed for *A. sagrei*, ocular globe morphogenesis in *C. calyptratus* includes pronounced asymmetric elongation along the lateromedial axis (Figure 6). In contrast with *A. sagrei, C. calyptratus* embryos exhibit comparable expansion along both the dorsoventral and nasotemporal axes (Figure 6). As with the anole, ocular elongation is followed by a period of retraction that ends when the globe regains a spherical shape.

**Figure 6.**
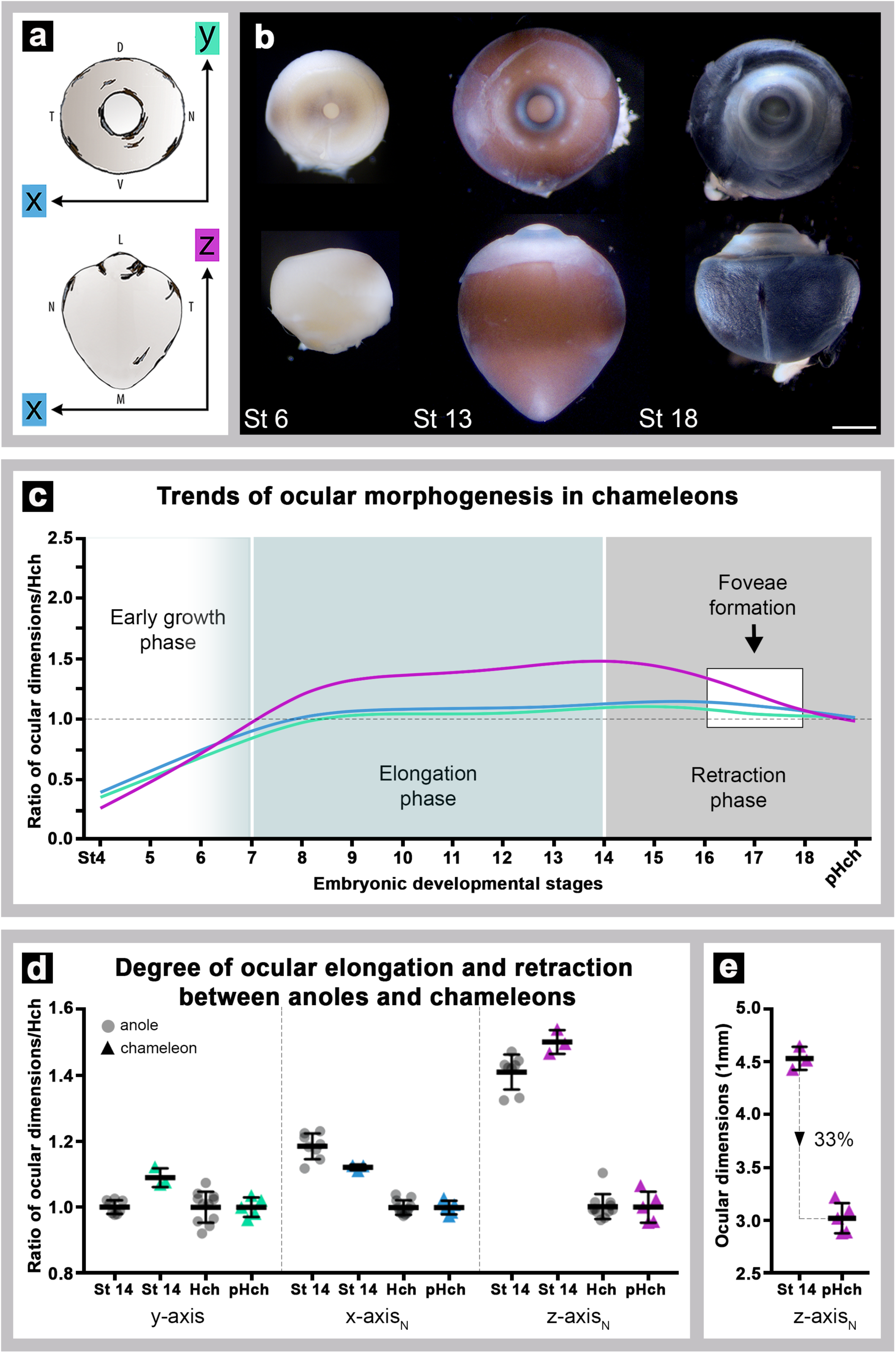
Chameleon eyes also undergo asymmetrical ocular elongation followed by retraction during embryonic development. Diagram (a) demonstrates how y- (green), x- (blue), and z-axis (magenta) ocular measurements were made. Image (b) shows lateral (top) and dorsal (bottom) views of whole fixed right eyes from early (St 6), mid (St 13), and late (St 18) embryonic development. Like the anole, ocular elongation (St 13) and retraction (St 18) phases are evident. All eyes are to scale; scale bar is 1 mm. Graph (c) summarizes trends in chameleon ocular morphogenesis throughout stages 4-pHch (pHch – just prior to hatching). This graph was generated by taking each lizard’s ocular dimensions (y-axis and normalized x- and z-axis_N_) and dividing it by the mean (µ) pHch ocular dimensions (y-, x-, and z-axis) to calculate a ratio. Graph (d) compares the degree of ocular elongation (St 14) and subsequent retraction (Hch and pHch) between anoles (grey circles) and chameleons (color triangles: y-axis – green, x-axis_N_ – blue, and z-axis_N_ – magenta). Calculations were made following same formula outlined above in graph (c) for each respective species. Graph (e) indicates the relative percentage of retraction occurring in chameleons between stage 14 and pHch.

Compared with the anole, the elongated region in chameleon is less acute and occurs over a broader area; in the anole, the elongated medial face appears more acute and funnel-like (Compare St 13 chameleon in Figure 6B to St 13 anole in Figure 4b). Despite this difference in morphology, the onset of ocular elongation and retraction timing is nearly identical to anoles. For instance, ocular elongation begins around stage 6 and peaks by stage 14. This is followed by ocular retraction, which starts at stage 15 and plateaus just before hatching (Figure 6c). The chameleon fovea also develops during the period of retraction (between stages 16-18) (Figure 6c), and its progression also matches that of the anole (data not shown).

## Discussion

The work we present here is a first step in establishing the brown anole as a new model organism for fovea developmental studies. A primary motivation for choosing this lizard is that the anole eye contains two fovea that represent the extremes of fovea morphology: a central fovea with a pit devoid of all retina cell layers and a temporal fovea with a shallow pit that retains the retinal layers. Most vertebrate species do not have fovea, but in those that do, the fovea can differ greatly in pit shape, depth, and diameter.^19,42-45^ In humans, only the GCL and INL retinal layers are laterally displaced, while the ONL is retained. This results in a broad and relatively shallow foveal pit with a high density of photoreceptor cells at its center.^30,46-50^ This is also true of most foveated non-human primates. Although in some species, the GCL and/or INL are retained, and as a result, only a rudimentary pit is present.^45,51-54^ The fovea of birds is variable, and, like the anole, some species are bifoveated, while others only possess a single fovea located either in the central or temporal retinal regions.^44^

Anoles are the only squamate genus known to have a bifoveated retina. Among the anole species studied, all have a prominent central fovea devoid of cell layers and a shallower temporal fovea that retains these layers.^19,20,55^ Slight variations in pit depth have been observed across anoles and correlate with prey size.^19^ For instance, anole species that routinely eat smaller prey have considerably deeper temporal foveae. As in birds, the location of the fovea in different squamate reptiles varies. In diurnal geckos only a temporal fovea is present.^56-60^ Although all layers are generally present in these lizards, the degree of lateral displacement ranges from partial to full layer retention. In *Lygodactylus*, which is a gekkonid, and *Gonatodes*, which is a sphaerodacylid, the GCL is nearly absent, while the INL is only thinned and ONL packing is present at the foveal pit center.^58^ In contrast, *Phelsuma* geckos have only the shallowest of depressions absent of any pronounced displacement of GCL and INL, resulting in a pit similar to the temporal fovea of anoles.^58,59^ In other lizards, including chameleons, the opposite is generally true, with most having a large prominent central fovea devoid completely of retinal cell layers.^61,62^

We observed four distinct phases of ocular morphogenesis in anoles and chameleons. The first was a period of symmetrical growth that occurred during the first week of embryonic development post-fertilization. At the end of this period, the retina was thicker and exhibited a mounded appearance in the prospective foveal regions (Figure 3g). This was followed by a second period, defined by asymmetrical growth, where the regions that eventually gave rise to the fovea became strikingly elongated. Coincidingly, this period also marks the gradually disappearance of retinal mounding within the foveal regions (*Rasys et. al*., in prep). By late development, when the foveae take on their characteristic morphology, these regions appear to undergo retraction coincident with retinal remodeling, i.e., pit formation and photoreceptor cell packing. The fourth phase was characterized by a uniform expansion of the globe. In both the brown anole and the veiled chameleon the regions that undergo elongation and retraction are localized to areas of the retina where the foveae develop. Additionally, these foveal regions, characterized by early retinal mounding, undergo retina differentiation and lamination prior to the rest of the retina (*Rasys et. al*., in prep). These observations suggest a relationship between changes in retinal differentiation, ocular shape, and foveal development.

We propose that ocular elongation followed by retraction are necessary steps in the retinal remodeling needed to generate a fovea in vertebrates. Consistent with this idea, evidence of asymmetrical globe development can be seen in the eyes of diurnal, but not nocturnal, gecko species. Examples of this phenomenon can be seen in Figures 2 and 3 of the publication by Guerra-Fuentes and colleagues addressing the embryology of the retinal pigmented epithelium in five species of sphaerodactyls,^32^ and in Figures 3 and 4 of the publication by Griffing, Gamble and Bauer addressing pigment development in nocturnal gekkonids (*Gekko kuhli, Lepidodactylus lugubris*) and diurnal gekkonids (*Phelsuma laticauda, Sphaerodactylus macrolepis*).^33^ Among mammals, asymmetric globe development has been observed only for foveated haplorrhine primates but not for non-foveated primates or other mammalian species.^24-31^ Together, these observations suggest that fovea morphogenesis is similar among foveated vertebrates. It will be interesting to learn if the eyes of foveated birds and fish undergo similar morphogenetic changes during development.

In the brown anole, we noted that the magnitude of asymmetric ocular shape changes during development appeared proportional to the extent of retinal remodeling associated with formation of the morphologically dramatic central fovea compared to the less distinct temporal fovea. We speculate that the process of elongation and retraction of required for fovea formation, and that the relative extent to which an eye elongates and retracts during fovea-genesis is directly proportional to the amount of retinal remodeling required to make that particular fovea. If true, this may explain the variability of fovea morphology present within each foveated species.

Several different mechanisms could mediate changes in the size and shape of the ocular globe during development. We hypothesize that changes in intraocular pressure (IOP) contribute to asymmetric ocular morphogenesis in the brown anole. This idea arises from our observations that the embryonic anole eye appears to be pressurized during peak periods of elongation and deflated during retraction. The idea that IOP can drive ocular growth is not a new one. Previous studies have shown that IOP plays a pivotal role in regulating normal ocular growth in chick^63^ and increases in IOP can lead to induced myopia (generalized axial elongation of the globe) in this animal.^64,65^ In foveated primates, Hendrickson and Springer proposed a model where high IOP induces pit formation due to inherent increased elasticity present at the foveal avascular zone, while “retinal stretching” induced by ocular growth, facilitates the centripetal movement of photoreceptor cells towards the foveal center.^66-68^ Although, it is possible that lack of blood vessels would predispose this region of the primate retina to be more susceptible to IOP and, therefore, form a pit, this cannot explain pit formation in the lizard. Anoles have a retina that is entirely avascular.^43,55^ This suggests that if regional differences in retinal elasticity are present in the anole, they are unlikely to be caused by avascular zones. Another challenge with this model, as applied to anoles, is that it requires IOP to be high for pit formation to occur. In the brown anole, pit formation occurs during the period that the eye is soft, indicating that IOP is low.

We propose, for anoles at least, that high IOP is involved in facilitating ocular elongation, but that IOP is comparatively low during ocular retraction and pit formation. As for Hendrickson and Springer’s model, there must be additional mechanisms that mediate regional differences in the elasticity of the foveal anlagen compared to other regions of the ocular globe. We think it likely that regional and dynamic changes in the elasticity of the tissues associated with the outer surface of the globe are required for normal eye development in foveated lizards, and may be true for primates as well.

## Supporting information

Supplementary Figures.docx

## Acknowledgements

The authors thank Aaron Alcala, Sergio Minchey, Sukhada Samudra, Christina Sabin, and Rebecca Ball of the Menke and Lauderdale research groups at the University of Georgia, and Diana Baumann, Richard Kupronis, David Jewell, Alex Muensch and Nikki Inlow of the Reptile and Aquatic Facility at the Stowers Institute for Medical Research, for their help with animal husbandry, maintenance and care of the anolis and chameleon colonies, respectively. The authors thank Drs. Jonathan Eggenschwiler, Heike Kroeger and Robert Hufnagel, and Christina Sabin, Sukhada Samudra, and Rida Osman for helpful discussions about this project and comments on the manuscript.

