## Supplementary Figures.docx for "Ocular elongation and retraction in foveated reptiles"

Supplementary Figure 1. Distribution of unaltered anole ocular measurements throughout embryogenesis. Graphs (a-c) show y- (green), x- (blue), and z-axis (magenta) ocular measurements between stage 4-Hch and adult lizards; markers indicate: Adt F – adult female; Adt M – adult male; and dashed line – mean (µ) Hch ocular dimension. Graph (d) summaries graphs (a-c).

Supplementary Figure 2. Normalizing ocular x- and z-axis ocular dimensions to the y-axis reduces variability in the dataset. Graphs (a-b) show the distribution of unaltered x- and z-axis (black) compared to the normalized values x- (blue) and z-axis_N_ (magenta). Calculations were made by taking each lizard’s x- or z-, dividing those values by its y-axis, then multiplying by the mean (µ) of the y-axis for its stage group.

Supplementary Figure 3. Chameleon unaltered ocular measurements throughout embryonic development. Graphs (a-c) show y- (green), x- (blue), and z-axis (magenta) ocular measurements between stage 4-pHch; dashed line represents mean (µ) pHch ocular dimension. Graph (d) summaries graphs (a-c).

Supplementary Table 1. Anole ocular axial dimensions across embryonic development and adulthood. Table shows the number of lizards collected from embryonic stages (St 4-Hch) and adults (Adt F – adult female; Adt M – adult male) as well as unaltered y-, x-, and z-axis and normalized (x- and z-axis_N_) ocular measurements; mean (µ) ± standard deviation and range are reported in microns.

Supplementary Table 2. Chameleon ocular dimensions throughout embryogenesis. Table shows the number of lizards collected days post oviposition (DPO) and their subsequent stages (St 4-pHch) following anole staging guidelines. Dataset includes unaltered y-, x-, and z-axis and normalized (x- and z-axis_N_) ocular measurements; mean (µ) ± standard deviation and range are reported in microns. Note: stages 5 and 9-11 have an “n” of 1; mean (µ) is reported as a single value and no range is provided.
