## Supplementary Figs Tables.pdf for "Ocular elongation and retraction in foveated reptiles"

### Supplemental Figure 1

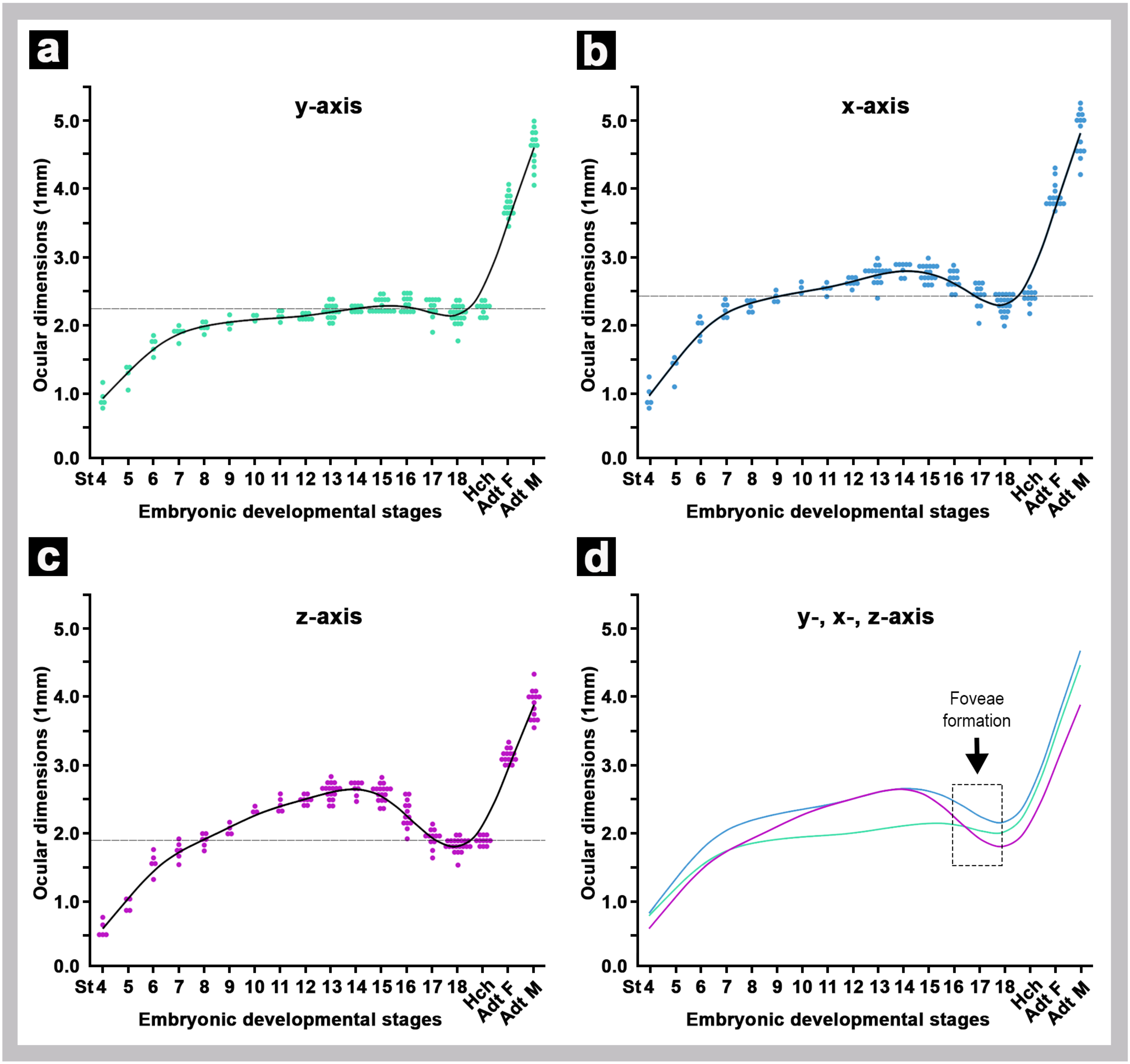

Supplemental Figure 2

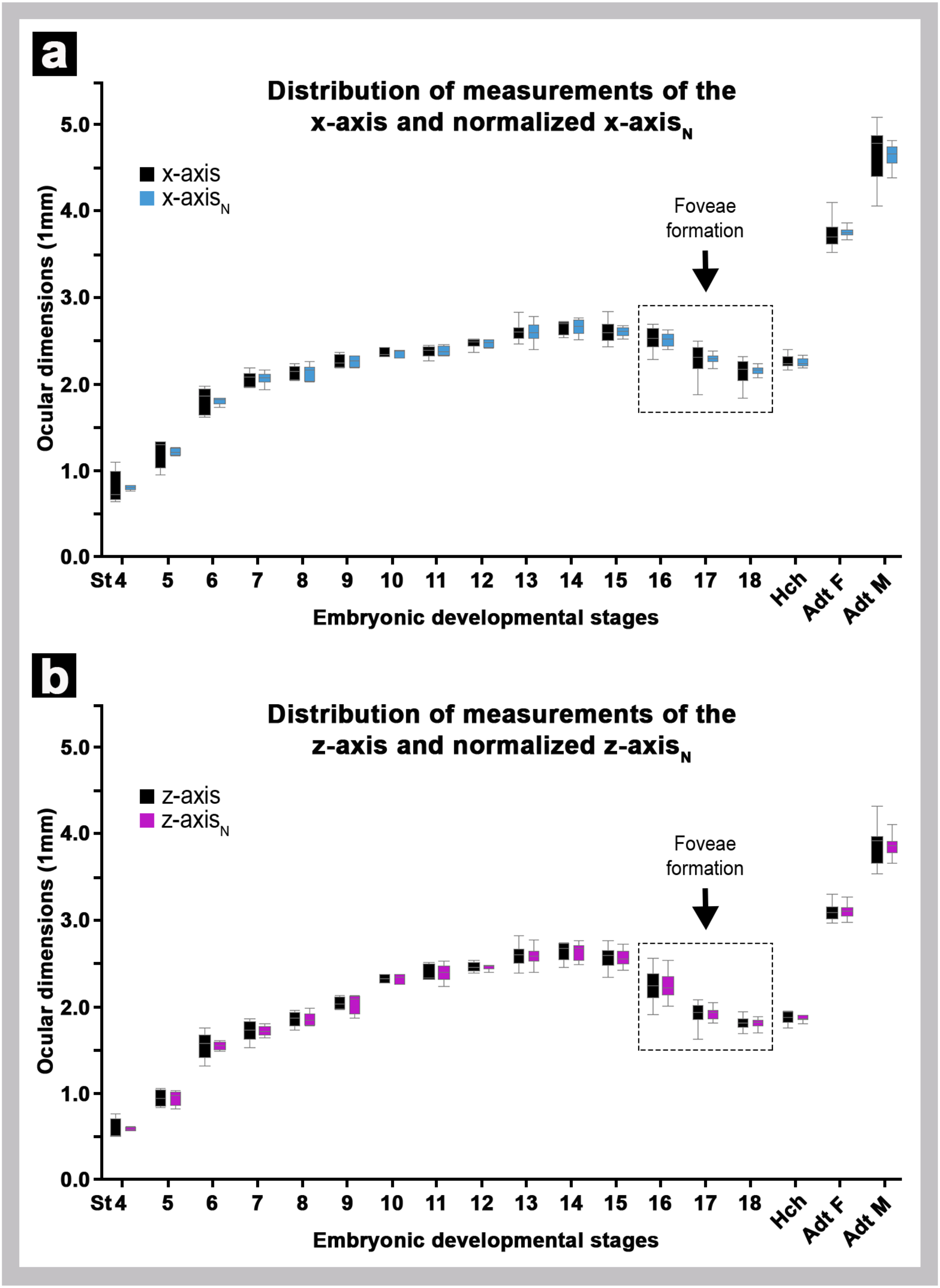

**Supplemental Figure 3**

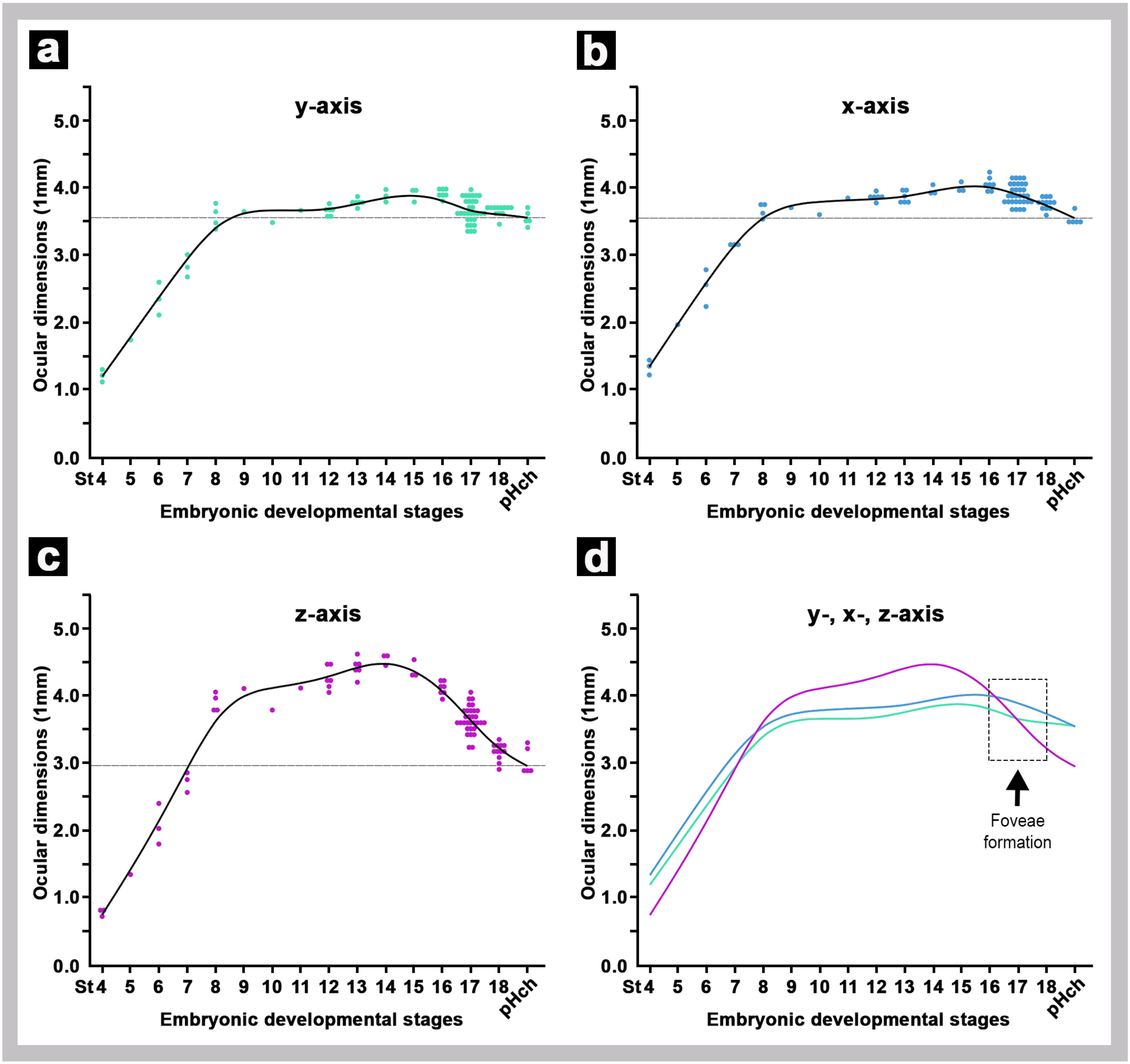

**Supplemental Table 1**

#### Anole ocular axial dimensions (μm)

| Stage | n | $\mu_y$ | $\mu_x$ | $\mu_z$ | $Range_y$ | $Range_x$ | $Range_z$ | $\mu_{x\,(norm)}$ | $\mu_{z\,(norm)}$ |
|---|---|---|---|---|---|---|---|---|---|
| 4 | 5 | 765 ± 155 | 808 ± 185 | 587 ± 116 | 640-1017 | 640-1098 | 502-762 | 805 ± 27 | 588 ± 20 |
| 5 | 4 | 1116 ± 142 | 1221 ± 181 | 947 ± 101 | 905-1203 | 953-1336 | 838-1059 | 1219 ± 44 | 952 ± 90 |
| 6 | 5 | 1564 ± 126 | 1808 ± 157 | 1552 ± 158 | 1383-1704 | 1619-1981 | 1316-1753 | 1808 ± 42 | 1550 ± 49 |
| 7 | 6 | 1749 ± 87 | 2066 ± 87 | 1722 ± 118 | 1586-1848 | 1959-2189 | 1532-1865 | 2068 ± 72 | 1722 ± 58 |
| 8 | 6 | 1818 ± 59 | 2137 ± 80 | 1859 ± 83 | 1717-1898 | 2043-2237 | 1735-1964 | 2138 ± 89 | 1860 ± 74 |
| 9 | 4 | 1884 ± 90 | 2262 ± 80 | 2042 ± 72 | 1797-2007 | 2188-2367 | 1973-2131 | 2263 ± 70 | 2045 ± 118 |
| 10 | 3 | 1952 ± 33 | 2362 ± 54 | 2327 ± 46 | 1914-1977 | 2321-2423 | 2280-2372 | 2362 ± 49 | 2327 ± 56 |
| 11 | 5 | 1963 ± 61 | 2382 ± 67 | 2393 ± 96 | 1896-2032 | 2273-2448 | 2315-2510 | 2383 ± 56 | 2394 ± 103 |
| 12 | 8 | 1974 ± 44 | 2474 ± 53 | 2460 ± 52 | 1917-2036 | 2371-2526 | 2393-2537 | 2475 ± 41 | 2460 ± 28 |
| 13 | 17 | 2050 ± 103 | 2590 ± 127 | 2585 ± 110 | 1875-2238 | 2250-2834 | 2389-2822 | 2592 ± 104 | 2588 ± 99 |
| 14 | 8 | 2091 ± 44 | 2662 ± 78 | 2646 ± 105 | 2041-2144 | 2536-2729 | 2457-2743 | 2663 ± 87 | 2647 ± 100 |
| 15 | 17 | 2132 ± 91 | 2610 ± 116 | 2568 ± 115 | 2031-2317 | 2435-2837 | 2345-2767 | 2611 ± 71 | 2569 ± 84 |
| 16 | 13 | 2153 ± 96 | 2515 ± 128 | 2254 ± 193 | 2016-2300 | 2289-2690 | 1911-2566 | 2515 ± 78 | 2253 ± 160 |
| 17 | 12 | 2101 ± 138 | 2290 ± 170 | 1918 ± 130 | 1751-2235 | 1881-2495 | 1631-2086 | 2290 ±58 | 1919 ± 64 |
| 18 | 21 | 2009 ± 136 | 2148 ± 130 | 1804 ± 92 | 1624-2242 | 1842-2323 | 1526-1947 | 2151 ± 68 | 1807 ± 62 |
| Hch | 11 | 2090 ± 98 | 2245 ± 98 | 1879 ± 62 | 1924-2244 | 2024-2403 | 1755-1955 | 2246 ± 50 | 1881 ± 71 |
| Adt F | 15 | 3576 ± 163 | 3739 ± 167 | 3103 ± 94 | 3301-3903 | 3524-4102 | 2971-3305 | 3740 ± 91 | 3105 ± 78 |
| Adt M | 14 | 4435 ± 267 | 4651 ± 298 | 3863 ± 212 | 3903-4782 | 4060-5082 | 3537-4321 | 4651 ± 116 | 3866 ± 128 |

**Supplemental Table 2**

#### Chameleon ocular axial dimensions (µm)

| Stage | DPO | n | $\mu_y$ | $\mu_x$ | $\mu_z$ | $Range_y$ | $Range_x$ | $Range_z$ | $\mu_{x\ (norm)}$ | $\mu_{z\ (norm)}$ |
|---|---|---|---|---|---|---|---|---|---|---|
| 4 | 90-96 | 3 | 1190 ± 79 | 1319 ± 102 | 771 ± 53 | 1106-1263 | 1208-1410 | 710-805 | 1319 ± 16 | 776 ± 100 |
| 5 | 96 | 1 | 1729 | 1954 | 1333 | - | - | - | 1954 | 1333 |
| 6 | 100 | 3 | 2340 ± 243 | 2515 ± 274 | 2063 ± 304 | 2099-2586 | 2225-2770 | 1785-2388 | 2514 ± 38 | 2057 ± 91 |
| 7 | 96-100 | 3 | 2823 ± 166 | 3143 ± 16 | 2712 ± 148 | 2665-2995 | 3128-3160 | 2551-2843 | 3150 ± 168 | 2712 ± 39 |
| 8 | 110-120 | 4 | 3558 ± 171 | 3634 ± 121 | 3873 ± 116 | 3374-3757 | 3521-3751 | 3754-3991 | 3636 ± 87 | 3876 ± 108 |
| 9 | 110 | 1 | 3636 | 3693 | 4094 | - | - | - | 3693 | 4094 |
| 10 | 110 | 1 | 3471 | 3588 | 3774 | - | - | - | 3588 | 3774 |
| 11 | 110 | 1 | 3653 | 3834 | 4101 | - | - | - | 3834 | 4101 |
| 12 | 104-120 | 6 | 3642 ± 68 | 3825 ±53 | 4232 ± 181 | 3556-3730 | 3763-3923 | 4036-4460 | 3826 ± 74 | 4232 ± 137 |
| 13 | 110-125 | 6 | 3749 ± 57 | 3824 ± 66 | 4406 ± 139 | 3682-3847 | 3744-3916 | 4187-4605 | 3824 ± 45 | 4405 ± 109 |
| 14 | 125 | 3 | 3847 ± 101 | 3950 ± 73 | 4532 ± 83 | 3778-3963 | 3895-4033 | 4437-4594 | 3951 ± 31 | 4533 ± 110 |
| 15 | 125-130 | 3 | 3892 ± 100 | 3990 ± 74 | 4371 ± 136 | 3777-3960 | 3947-4075 | 4262-4523 | 3991 ± 99 | 4371 ± 109 |
| 16 | 130-135 | 7 | 3875 ± 72 | 4040 ± 92 | 4085 ± 90 | 3792-3971 | 3912-4177 | 3935-4179 | 4040 ± 42 | 4085 ± 56 |
| 17 | 127-163 | 31 | 3620 ± 170 | 3873 ± 142 | 3622 ± 188 | 3328-3898 | 3644-4139 | 3201-4038 | 3876 ± 122 | 3623 ± 117 |
| 18 | 163-171 | 12 | 3629 ± 65 | 3732 ± 72 | 3142 ± 118 | 3447-3690 | 3580-3831 | 2893-3274 | 3733 ± 59 | 3142 ± 115 |
| pHch | 190-195 | 5 | 3532 ± 105 | 3521 ± 91 | 3023 ± 207 | 3398-3657 | 3460-3682 | 2827-3287 | 3522 ± 73 | 3021 ± 142 |

### *Anolis sagrei* eye development

Ashley M. Rasys[1,2], Shana H. Pau[3], Katie E. Irwin[1], Sherry Luo[3], Douglas B. Menke[3], and James D. Lauderdale[1,4]
[1]Department of Cellular Biology, [2]College of Veterinary Medicine, [3]Department of Genetics, [4]Neuroscience Division of the Biomedical and Health Sciences Institute,
The University of Georgia, Athens, GA 30602, USA

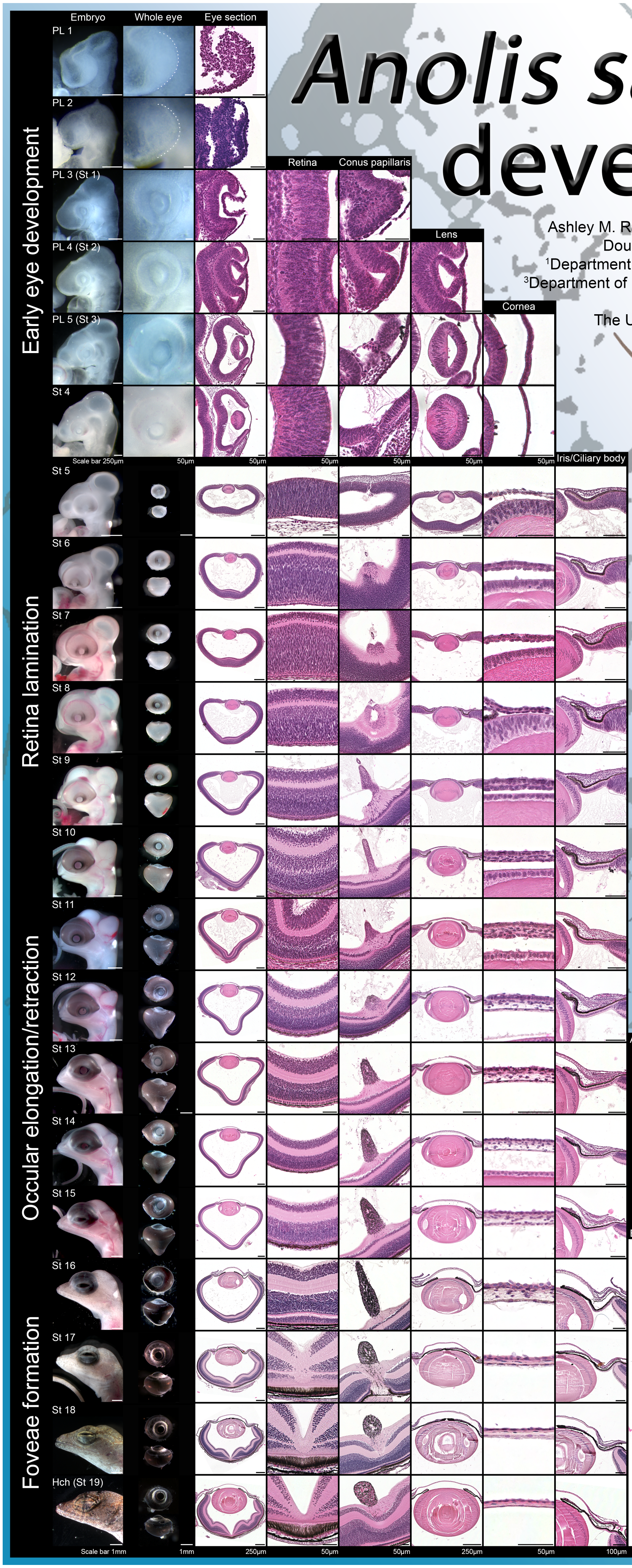


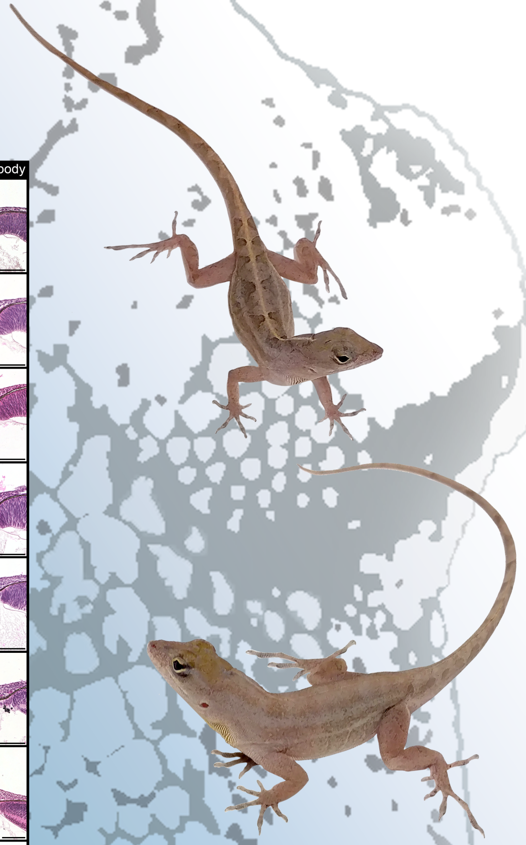

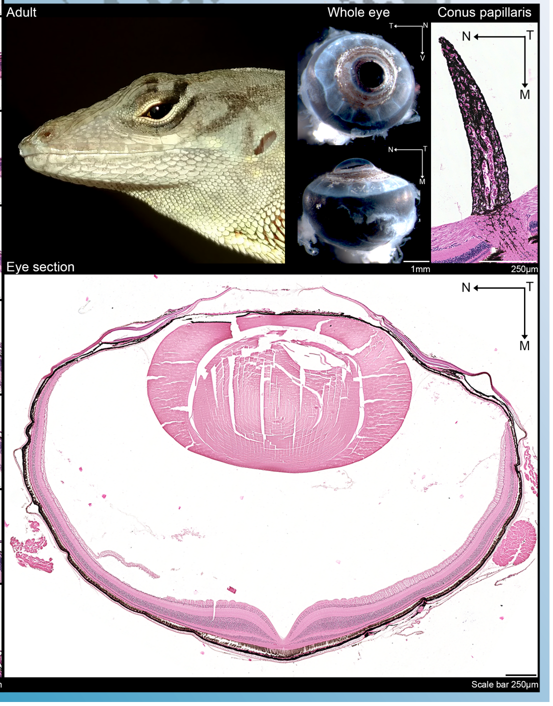
